# Cerebellar Purkinje cell activity modulates aggressive behavior

**DOI:** 10.1101/2020.01.06.891127

**Authors:** Skyler L. Jackman, Christopher H. Chen, Heather L. Offermann, Iain R. Drew, Bailey M. Harrison, Anna M. Bowman, Katelyn M. Flick, Isabella Flaquer, Wade G. Regehr

## Abstract

Although the cerebellum is traditionally associated with balance and motor function, it also plays wider roles in affective and cognitive behaviors. Evidence suggests that the cerebellar vermis may regulate aggressive behavior, though the cerebellar circuits and patterns of activity that influence aggression remain unclear. We used optogenetic methods to bidirectionally modulate the activity of spatially-delineated cerebellar Purkinje cells to evaluate the impact on aggression in mice. Increasing Purkinje cell activity in the vermis significantly reduced the frequency of attacks in a resident-intruder assay. Reduced aggression was not a consequence of impaired motor function, because optogenetic stimulation did not alter motor performance. In complementary experiments, optogenetic inhibition of Purkinje cells in the vermis increased the frequency of attacks. These results establish Purkinje cell activity in the cerebellar vermis regulates aggression, and further support the importance of the cerebellum in driving affective behaviors that could contribute to neurological disorders.

## Introduction

Profound motor deficits such as ataxia and loss of oculomotor control are the most obvious manifestation of cerebellar damage. This has contributed to the popular view that the cerebellum is involved primarily in motor function, but this is far from a complete view of the behavioral functions of the cerebellum. fMRI studies suggest that some regions of the cerebellar cortex are devoted to motor function, but other regions are involved in working memory, language, emotion, executive function and many other nonmotor functions (Stoodley & Schmahmann, 2009; Van Overwalle *et al*., 2014). The cerebellum is also implicated in autism spectrum disorder (Wang *et al*., 2014), anxiety (Moreno-Rius, 2018), attention deficit disorder (Berquin *et al*., 1998), schizophrenia (Andreasen & Pierson, 2008), and other nonmotor neurological disorders (Phillips *et al*., 2015).

The posterior vermis region of the cerebellar cortex is particularly intriguing with regard to involvement in nonmotor behaviors. Damage to the cerebellar vermis in adults can lead to deficits in executive function, spatial cognition, linguistic processing, affect regulation, irritability, anger, aggression, and pathological crying or laughing (Schmahmann & Sherman, 1998; Levisohn *et al*., 2000). There is also extensive evidence suggesting that the vermis influences aggression. In seminal studies mapping the somatotopic organization of the cerebellar cortex, the Italian physiologist Guisseppe Pagano found that injecting curare into the vermis caused the animal to “become suddenly furious, and throw itself at those present, trying to bite them” or to “jump into the air, struggling to bite who knows how many phantoms of its agitated psyche” (Pagano, 1904). Later lesions studies demonstrated that resection of the vermis had the opposite influence on behavior and produced a calming effect (Sprague & Chambers, 1959; Peters & Monjan, 1971; Berman *et al*., 1974). Electrical stimulation of the deep cerebellar nuclei has been shown to drive aggressive behaviors such as sham rage (Zanchetti & Zoccolini, 1954) and attack (Reis *et al*., 1973). In human clinical studies, stimulating the surface of the vermis improved emotional control and reduced aggressive outbursts (Heath, 1977), and reduced feelings of anger (Cooper *et al*., 1976).

While previous research has implicated the vermis in regulating aggression, these studies have been largely anecdotal and did not define the cerebellar circuits that modulate aggression. Purkinje cells (PCs), the sole output cells of the cerebellar cortex, fire continuously at up to 100 Hz, and inhibit neurons in the deep cerebellar nuclei (DCN) that in turn influence other brain regions. Electrical stimulation of the cerebellar cortex activates all types of neurons in the vicinity of the electrode, including PCs. Molecular layer interneurons will also be activated, and they inhibit PC firing. Stimulation also antidromically activates mossy fibers, climbing fibers, and modulatory inputs from other regions, which might contribute to the behavioral consequences of stimulation. For these reasons it is difficult to know how electrical stimulation of the vermis influences behavior. For lesion studies of the cerebellar vermis, it is not clear how the firing properties of downstream DCN neurons were altered. Thus, prior studies of the cerebellum and aggression are difficult to interpret.

To determine the role of cerebellar outputs in regulating aggression, we used optogenetic techniques to selectively control PC activity in mice. We determined the effect on behavior using the resident-intruder assay, a measure of natural territorial aggression in rodents. Manipulating PC firing in the vermis, but not in other cerebellar regions, enabled rapid, bidirectional control of aggression. This study establishes that in the cerebellar vermis elevated PC firing suppresses aggression, whereas suppressing PC firing promotes aggressive behavior.

## Results

To selectively modulate the activity of PCs in the cerebellar vermis, we used PCP2-cre mice (Zhang *et al*., 2004) to restrict expression of the microbial opsins ChR2 (channelrhodopsin-2, (Boyden *et al*., 2005)) and NpHR3.0 (halorhodopsin, (Zhang *et al*., 2007)) to PCs. *In vitro* electrophysiological recordings (Figure 1a) confirmed that ChR2 stimulation could drive graded increases in PC firing that scaled with light intensity (Figure 1b), as previously reported (Guo *et al*., 2016). Similarly, halorhodopsin stimulation decreased PC firing rates in a light intensity-dependent manner (Figure 1c). To manipulate PC activity *in vivo*, optical fibers were chronically implanted in adult (>P42) male PCP2-cre::ChR2 or PCP2-cre::NpHR mice over the surface of the cerebellar vermis. Fibers were positioned at the midline over lobule VII (Figure 1 – Supplement 1), a region suggested to play a role in emotional processing (Stoodley & Schmahmann, 2009). Subsequent *in vivo* recordings through an adjacent craniotomy confirmed the ability of light to reliably increase or decrease the firing rate of putative PCs expressing ChR2 or halorhodopsin, respectively (Figure 1d-f).

**Figure 1.**
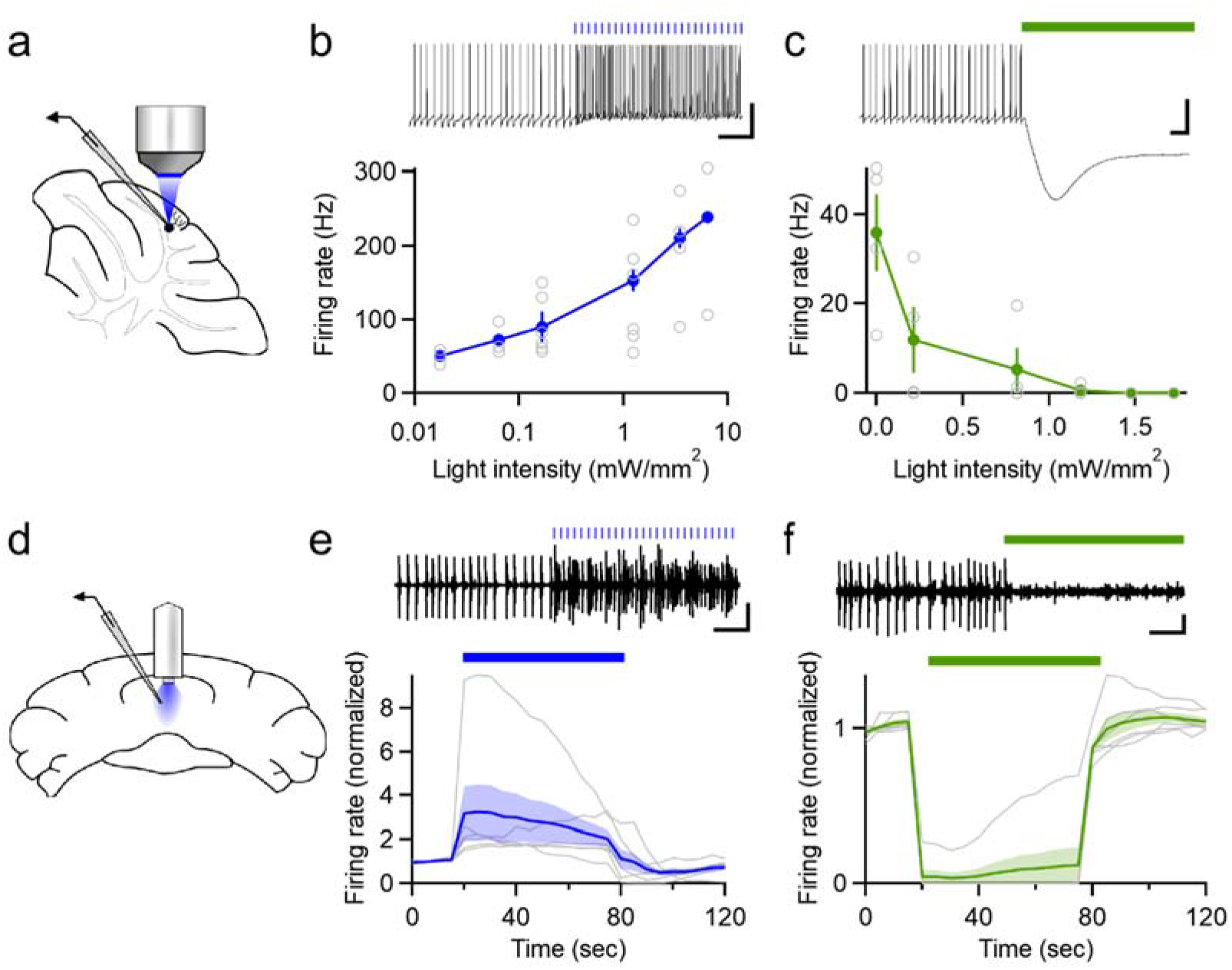
Optogenetic control of Purkinje cell activity. (**a**) Recording schematic for *in vitro* recording and optogenetic stimulation. (**b**) Firing rates elicited by ChR2 stimulation at different intensities (0.5 ms flashes, 50 Hz, n = 5). (**c**) Inhibition of PCs at different light intensities (sustained illumination, n = 4). (**d**) Schematic for recording PC activity during *in vivo* stimulation through a chronic fiber optic implant. (**e**) Top: Representative single unit recording during ChR2 stimulation and (bottom) average firing rate (n = 6). (**f**) Single unit recordings during halorhodopsin stimulation (n = 6). Average data in all figures represents mean ± SEM.

After allowing at least 1 week for recovery from fiber implantation surgeries, animals were placed in an open field arena and stimulated with increasing light intensities to determine if manipulating PC activity produced overt motor deficits or behavioral consequences. In ChR2-expressing mice, strong vermal stimulation using the highest light intensities (~110 mW/mm^2^ at the face of the fiber optic implant) often resulted in clear motor effects, causing mice to become immobile or exhibit seizure-like and dystonic activity. This behavior resembled previous descriptions of seizure-like activity driven by strong electrical stimulation of the vermal cortex (Chambers, 1947). Thus, for all subsequent assays we tailored the intensity of light delivered to each animal to the maximal intensity where animals remained mobile in the open field arena and did not display signs of motor impairment. In contrast, halorhodopsin-expressing animals exhibited no obvious behavioral effects in response to stimulation at the maximal light intensity deliverable by the fiber-coupled LED light source (61 mW/mm^2^). This value was used for all subsequent assays.

Because the cerebellum plays well-established roles in motor control and balance, we first tested whether manipulating vermal PC firing caused more subtle motor impairments than those described above, that might interfere with the expression of other behaviors. To evaluate coordination, animals were tested on accelerating rotarod assays for two consecutive trials during which they received either optical stimulation or no stimulation. Neither excitation with ChR2 (Figure 2a) nor inhibition with halorhodopsin (Figure 2b) affected the rotarod performance. We next assessed locomotion during open-field assays. Animals received alternating 3-minute blocks of optogenetic excitation (Figure 2c). Automated animal tracking (Figure 2d) revealed that optogenetic activation of PCs had no effect on distance traveled, nor the time animals spent in the center of the arena, a measure of anxiety (Figure 2e). To assess the effect of stimulation on locomotion with greater temporal resolution, we averaged animal speed across all blocks of stimulation within the trial, centered around the onset of stimulation, and found that stimulation did not induce any transient change in locomotion (Figure 2 – figure supplement 1). Similarly, optogenetic inhibition did not affect locomotion or anxiety (Figures 2f-g and Figure 2 – figure supplement 1). Together, these data suggest that manipulating vermal PC firing does not strongly affect coordination, locomotion or anxiety.

**Figure 2.**
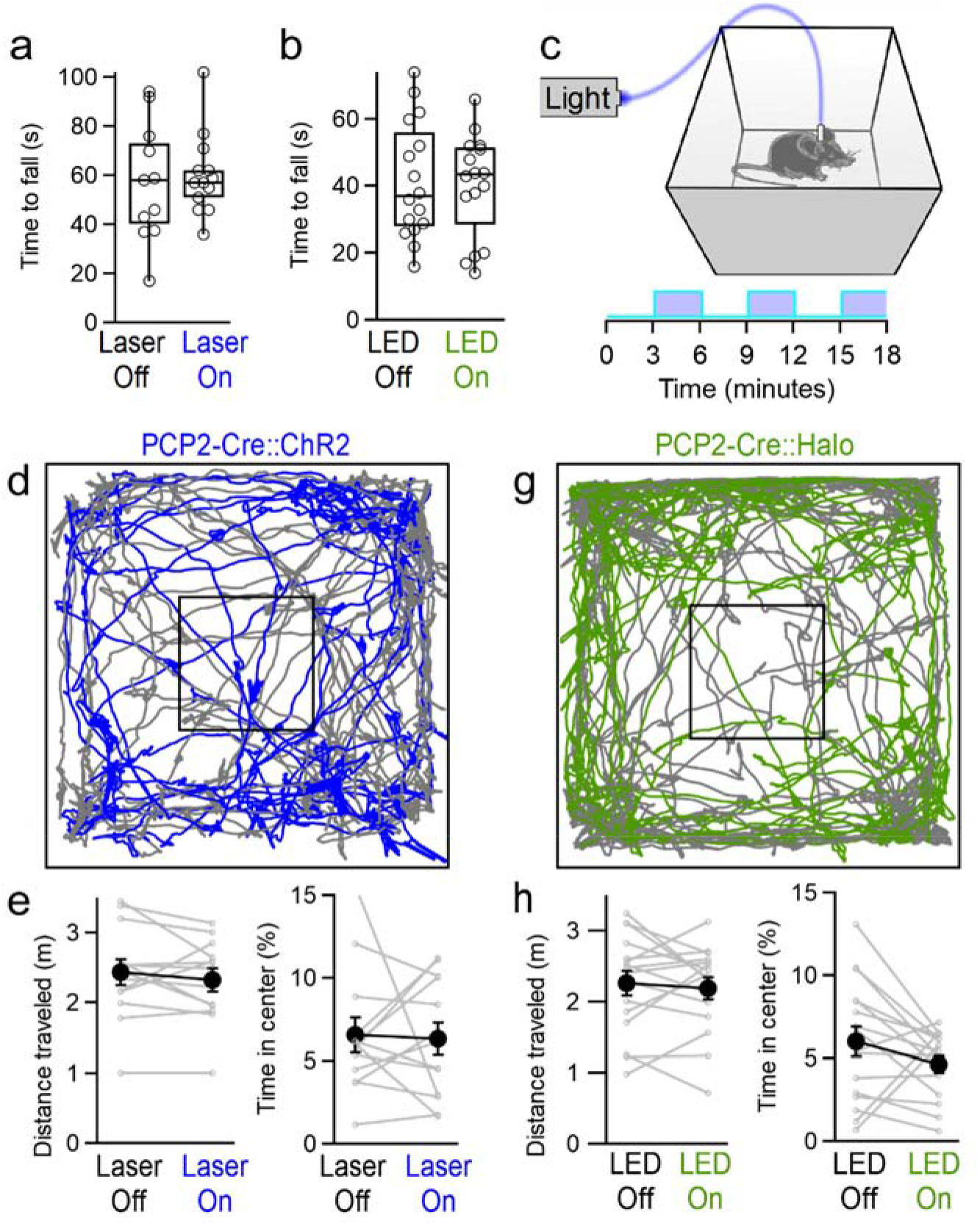
Manipulating vermal Purkinje cell activity does not affect coordination, locomotion or anxiety. (**a**) Time to fall for rotarod assays during ChR2-mediated excitation. Mice were tested in 2 consecutive trials, and randomly assigned to receive stimulation during either the first or second trial. (n = 13) (**b**) Same as in (a), but for halorhodopsin-mediated inhibition of vermal PC firing. (n = 16) (**c**) Schematic for open field assay with optogenetic stimulation. (**d**) Representative tracking data throughout alternating periods with stimulation (blue) and without (gray). (**e**) Total distance traveled and time spent in the center of the arena for mice during epochs with and without stimulation of vermal PCs (n = 13). G-H, Same as D-E but for halorhodopsin-mediated inhibition of vermal PCs (n = 17).

To assess the impact of cerebellar activity on social and aggressive behaviors we performed resident-intruder assays while optogenetically manipulating PC activity in the resident (aggressor) animal (Figure 3a). Although resident mice reliably display aggressive behaviors in resident-intruder assays, attacks occur at a relatively infrequent rate of <1 attack per minute (Leypold *et al*., 2002; Yang *et al*., 2013; Lewis *et al*., 2015). In order to increase baseline aggression, fiber-implanted mice were housed with females, providing the opportunity to mate, then subsequently singly-housed for at least 1 week prior to assays. Adult male BALB/c intruders were introduced into the resident’s home cage for 10 minutes. Optogenetic stimulation was delivered to residents in alternating 1-minute blocks. The onset and duration of multiple behaviors were recorded, including aggression (attacks, tail rattles, chasing and lateral threat), social encounters (face-to-face contact and ano-genital sniffing), as well as self-grooming by the resident (Figure 3b).

**Figure 3.**
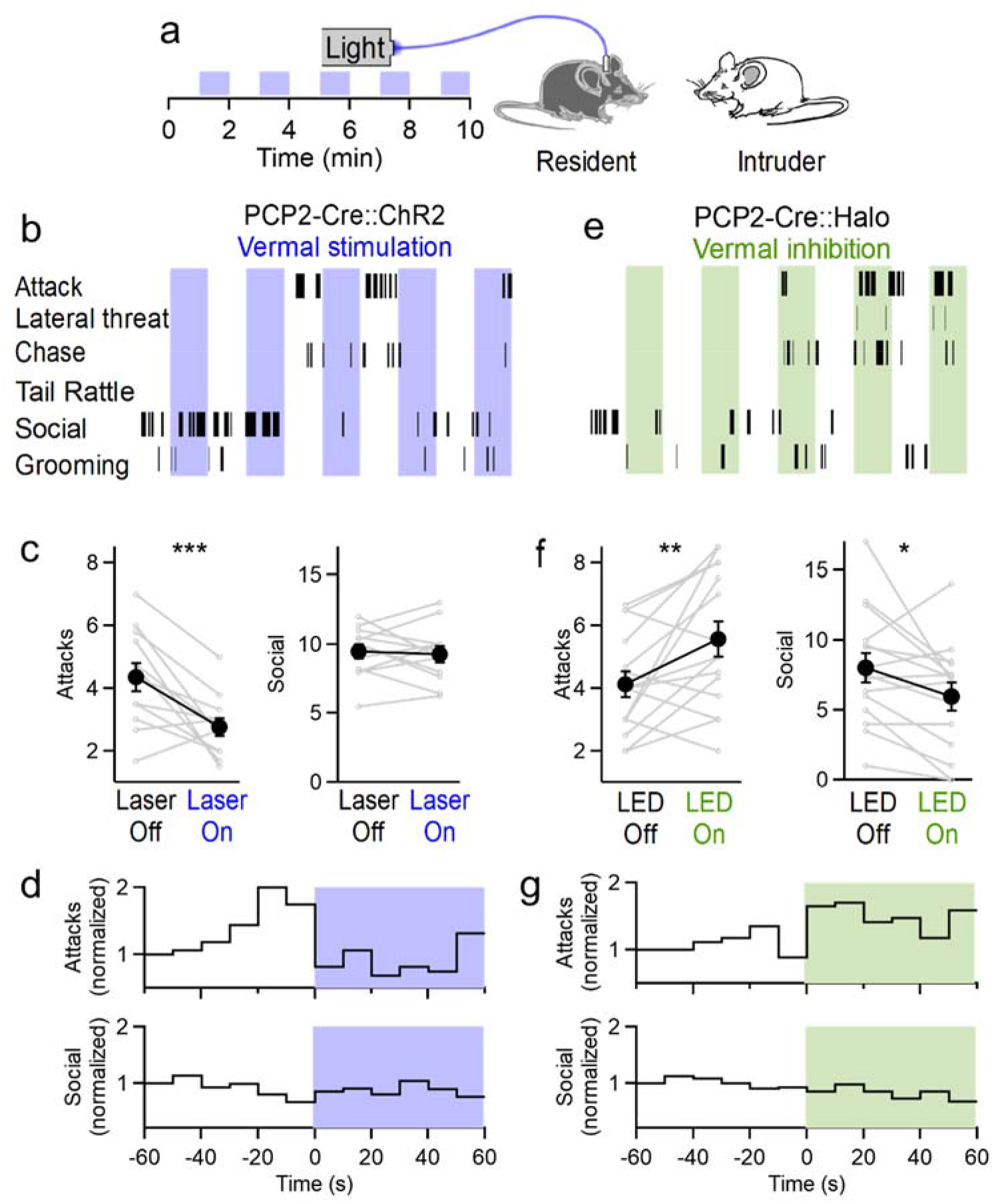
Bidirectional control of aggression by optogenetic modulation of vermal Purkinje cell activity. (**a**) Schematic for resident-intruder assays with optogenetic stimulation. (**b**) Representative scoring of social and aggressive behaviors. (**c**) Average number of attacks and social encounters during ChR2 assays (31 assays from 12 residents). (**d**) Peristimulus time histogram of normalized frequency of attacks (top) and social investigations (bottom) during epochs with and without ChR2-mediated excitation of vermal PCs. (**e-g**), Same as in (b-d), but during Halorhodopsin-mediated inhibition of vermal PCs (34 assays from 15 residents).

Optogenetic activation of vermal PCs significantly decreased the number of attacks (*p* = 0.003, *t*-test) (Figure 3c). Stimulation did not affect the frequency of social interactions (*p* = 0.7, *t*-test), or the rate of tail rattles, chasing, or lateral threat, though it did increase the rate of self-grooming by the resident (Figure 3 -figure supplement 1). An advantage of the optogenetic approach we have used is that it allows us to precisely determine the time course of the effect of stimulation on aggressive behavior with greater temporal resolution. We binned attacks in 10 second increments and averaged across alternating blocks at the onset of stimulation. Even though attacks are infrequent and stochastic, this analysis revealed that optical activation of PCs immediately reduced attack frequency, and when illumination was stopped the attack frequency gradually ramped up in the subsequent minute (Figure 3d). Stimulation reduced the frequency of attacks by 54% in the 10 seconds immediately following the onset of stimulation (Figure 3d). To put this into context, genetically ablating neurons in the ventromedial hypothalamus, a brain region colloquially referred to as the “attack area” because of its importance in regulating aggression, decreases the attack frequency by a little more than 50% (Yang *et al*., 2013).

To test whether decreased attacks might result from a distracting influence of light escaping from the implanted optical fiber, we performed resident-intruder assays with a separate cohort of wildtype mice that did not express opsins, and found that optical stimulation had no effect on either attacks or social interactions (Figure 3 – figure supplement 2). To test whether the effect on aggression was specific to stimulating activity in the vermis, we repeated the experiments in ChR2-expressing animals but implanted the optical fiber over Crus II, a region that has not been implicated in regulating aggression. In these mice, stimulating PC firing had no effect on the frequency of attacks (Figure 3 – figure supplement 2). Together, these results suggest that increased PC firing in the cerebellar vermis results in a rapid and significant decrease in aggression.

If elevating PC firing in the vermis decreases aggression, does suppressing PC firing increase aggression? It is not possible to address this question with electrical stimulation, but it possible using optogenetics. Inhibiting PCs with halorhodopsin (Figure 3e) had opposing effects on aggression, significantly increasing the number of attacks (*p* = 0.01, *t*-test), and decreasing social interactions (*p* = 0.03) (Figure 3f). Averaging the attack frequency across multiple epochs of stimulation showed that attack frequency nearly doubled in the 10 seconds following the onset of halorhodopsin-driven inhibition of PCs (Figure 3g). These results indicate that the activity of PCs in the cerebellar vermis exerts a bidirectional influence over aggressive behavior.

## Discussion

Here we demonstrate that Purkinje cell activity in the posterior vermis drives rapid, bidirectional changes in aggressive behavior. Several aspects of this present study provide important advances over previous studies that implicated the cerebellum in the regulation of aggression. First, we established that cerebellar activity regulates rodent aggression in an established assay that is amenable to quantification. This approach opens the door for quantitative studies in a genetically-manipulatable animal model, and promises to be beneficial for future studies of cerebellar control of aggression. Second, given the role of the cerebellum in motor control, it was important to establish that effects on aggression were not a secondary consequence of an alteration in motor control or motor performance. We evaluated this using open field and rotorod assays, and found that the same stimulation that altered aggression did not affect motor performance. Previous studies did not perform such a quantitative evaluation of motor performance. Third, the stimulation we used to suppress behavior was more selective than could be achieved with the electrical stimulation employed in previous studies, which in addition to stimulating PCs directly, can activate modulatory fibers, mossy fibers, climbing fibers, and inhibitory neurons in the cerebellar cortex. Consequently, in our ChR2 experiments, we can attribute decreased aggression to an increase in PC activity. Finally, our ability to reversibly increase aggression by suppressing PC firing indicates that the cerebellum can rapidly and bidirectionally regulate aggression. Previously, it was difficult to interpret the effects of lesions and irreversible damage to the cerebellum (Berman *et al*., 1974).

The present study raises a number of important questions regarding the manner in which the cerebellum controls behavior. What specific region of the cerebellar cortex is involved? We find that manipulating the activity of Purkinje cells in the posterior vermis is sufficient to significantly modulate aggression. This is consistent with clinical studies implicating lobule VII of the vermis in affective processing (Stoodley & Schmahmann, 2009). More detailed studies that manipulate activity in other areas of the midline vermis could add clarity to the specific regions of the cerebellar cortex that regulate aggression. It is also possible that more specific regulation of PC activity in the region controlling aggression (for example, without affecting PC firing in neighboring regions that alter other behaviors) will lead to larger effects on aggression. Furthermore, what is the nature of inputs that control this region? Different regions of the cerebellar cortex typically combine mossy fiber inputs from diverse sources, and it will be interesting to determine how these inputs are combined within the cerebellum to control aggression. Finally, what is the output pathway and the downstream targets that are ultimately regulated by activity in this region of the cerebellar cortex? Anatomical studies have described connections between the cerebellum and regions implicated in aggression, including hypothalamus (Haines *et al*., 1997) and prefrontal cortex (Kelly & Strick, 2003; Suzuki *et al*., 2012). Electrophysiological recordings have found that cerebellar stimulation evokes responses in those regions, along with limbic structures such as the hypothalamus, amygdala, and hippocampus (Anand *et al*., 1959; Snider & Maiti, 1976). Yet, while the somatotopic organization of the cerebellum is well characterized in regions that influence motor function, the output pathways of areas like the posterior vermis have yet to be clearly defined.

It is interesting to speculate on the nature of the role of the cerebellum in controlling aggressive behavior. The cerebellum has expanded in size relative to the cerebral cortex over the course of human evolution (Weaver, 2005), it contains more than half the neurons in brain and it possesses myriad connections to other brain regions. It is unsurprising that its influence should extend beyond the motor realm. Experiments on motor control suggest that the cerebellum combines inputs to generate predictions. It is natural to think that this computational strategy might be used by the posterior vermis of the cerebellum to learn how to respond to cues, and to ultimately decide when aggression is the correct response. Perhaps even subtle dysfunctions or misdirected plasticity within this region can lead to inappropriate aggressive behavior. For example, cerebellar damage often occurs in patients with PTSD (Rabellino *et al*., 2018). As non-invasive stimulation techniques like transcranial magnetic stimulation of the cerebellum emerge as a clinical treatment options (Demirtas-Tatlidede *et al*., 2010), it is increasingly important to understand the which areas of the cerebellum control nonmotor behaviors (Kelly & Strick, 2003). Future work could shed light on the anatomical projections and physiological impact of non-motor regions of the cerebellum.

## Materials and methods

### Animals

All experiments were conducted in accordance with federal guidelines and protocols approved by the Harvard Medical Area Standing Committee on Animals. Male mice of the following strains were used: Resident mice were either wild-type (WT) C57BL/6N (Charles River Laboratories), or Pcp2-cre mice (Jackson Laboratory, stock number 010536) crossed to either ChR2-EYFP (Ai32, Jackson Laboratory, 024109) or eNpHR3.0-EYFP (Halo) mice (Ai39 Jackson Laboratory, 014539). Intruder mice were BALB/c (Charles River Laboratories). YFP fluorescence in Ai32 and Ai39 mice was imaged using a Zeiss Axio Imager or Olympus MVX10 Macro dissecting microscope, and images were contrast enhanced in Fiji for visualization.

### *In Vitro* Physiology

Sagittal cerebellar slices were prepared from adult mice (P30-P100) and recordings were performed as previously described (Jackman *et al*., 2014). Briefly, animals were anesthetized with isoflurane and euthanized by decapitation. Brains were removed into oxygenated ice-cold cutting solution containing (in mm): 82.7 NaCl, 65 sucrose, 23.8 NaHCO_3_, 23.7 glucose, 6.8 MgCl_2_, 2.4 KCl, 1.4 NaH_2_PO_4_, and 0.5 CaCl_2_. Sagittal slices from the cerebellar vermis (250 μm thick) were prepared in ice-cold cutting solution using a Leica VT1200s vibrotome. Slices were transferred for 30 min into oxygenated artificial CSF (ACSF) at 32°C containing the following (in mm): 125 NaCl, 26 NaHCO_3_, 25 glucose, 2.5 KCl, 2 CaCl2, 1.25 NaH_2_PO_4_, and 1 MgCl_2_, adjusted to 315 mOsm, and allowed to equilibrate to room temperature for >30 min prior to recording. PCP2-Cre::ChR2-EYFP were used for all ChR2 recordings. Halorhodopsin recordings were performed in PCP2-Cre mice where opsin expression was driven by stereotaxic cerebellar injections (as previously described (Jackman *et al*., 2014)) of AAV9.EF1a.DIO.eNpHR3.0-eYFP.WPRE.hGH (Addgene26966). Although these mice were not used for behavioral experiments, similar optical sensitivity was observed in recordings performed for a separate study using PCP2-Cre::Ai39 mice (Guo *et al*., 2016).

Data were acquired using a Multiclamp 700B amplifier (Molecular Devices) digitized at 10 kHz with an ITC-18 (Instrutech), and low-pass filtered at 4 kHz. Acquisition and analysis were performed with custom software written in IgorPro (generously provided by Matthew Xu-Friedman, SUNY Buffalo). Whole-cell current clamp or on-cell recordings were obtained using borosilicate patch pipettes (2–4 MΩ), the internal solution contained the following (in mm): 150 K-gluconate, 3 KCl, 10 HEPES, 0.5 EGTA, 3 MgATP, 0.5 GTP, 5 phosphocreatine-tris2, and 5 phosphocreatine-Na2, with the pH adjusted to 7.2 with NaOH. Optical stimulation was delivered through the excitation pathway of a BX51WI upright microscope (Olympus) by either a 50 mW DPSS analog-controllable 473 nm blue laser (MBL-III-473-50mW, Optoengine), or a 590 nm Amber LED (160 mW, ThorLabs).

### Chronic fiber implantation and *in vivo* stimulation

Optical fiber implants were assembled as previously described (Sparta *et al*., 2011). Briefly, a multimode optical fiber (Thorlabs, NA 0.39, 200 μm core) was secured into ceramic ferrules (Thorlabs, 1.25 mm O.D.) with epoxy. Fibers were cleaved to protrude 0.2 mm below the ferrule, and the connector end was polished. Only fibers with >70% transmissivity were used. To determine the intensity of light exiting fibers, the output of fibers was measured with a power meter (Ophir; Vega). A photodiode was used to measure the relative intensity during short flashes controlled by the analog trigger of the laser, and this value was used to compute the power output during short flashes. The intensity of light delivered in vivo was computed by dividing the total light output (4.1 mW for the 473 nm laser, 2.3 mW for the 590 nm LED) by the surface area of the optical fiber. Optical fibers were implanted as described previously (Sparta *et al*., 2011). Briefly, adult mice (P40–P80) were anaesthetized with ketamine/xylazine (100/10 mg/kg) supplemented with isoflurane (1%–4%). An incision was performed to expose the skull, and the connective tissue and musculature above the cerebellum was gently peeled back. For vermal implants, the site for the craniotomy was determined using a fine pipette attached to a stereotaxic device (Kopf). After locating bregma, the pipette was moved caudal to the cerebellum, lowered 2.0 mm relative to bregma, then advanced rostrally until it touched the surface of the exposed skull. The site of CrusII craniotomies were determined similarly, but 1.5 mm ventral and 2.5 mm lateral of bregma. A craniotomy was performed at this site, and implants were lowered into place. Implants were secured to the skull using Metabond (Parkell), and the wound was sutured. Buprenorphine (0.05 mg/kg) was postoperatively administered subcutaneously every 12 hr for 48 hr.

### *In Vivo* Physiology

Mice from behavioral experiments were heavily anesthetized with isoflurane (2%). Anesthesia was maintained for all following procedures. A craniotomy immediately lateral to the implanted optical fiber was made to insert an electrode for extracellular recordings. A headplate was cemented (Metabond) anterior to the optical fiber, and the mouse was head-fixed for recordings. Electrodes were pulled from borosilicate glass (Sutter), filled with ACSF, and were inserted at an angle between 20 and 45 degrees to record single unit activity below the optical fiber. Most neurons were recorded between 1 and 2 mm from this entry point. Signals were acquired at 20 kHz between 0.2 and 7.5 kHz (Intan Technologies). Purkinje cells were identified by the presence of complex spikes, characteristic increase in noise as the electrode entered the Purkinje cell layer, and/or responsiveness to light. Single units were the sorted offline in Offline Sorter (Plexon) and analyzed in Matlab (Mathworks).

### Behavior

Mice used in behavioral experiments were housed in a 12 hour reverse light-dark cycle (lit 7PM-7AM). The timeline for experiments were as follows: Resident (aggressor) mice were allowed to recover from implants surgeries for at least 7 days. They were then paired with an adult C57BL/6N female for 7-12 days. The female was removed and the resident mouse remained in social isolation for at least 7 days. No cage changes were performed during social isolation to enhance subsequent territorial dominance aggression. Residents were first tested for signs of stimulation-induced motor disfunction, then assayed in the open field, rotarod, and finally aggression (resident-intruder) over the course of several weeks. Prior to behavioral experiments, animals were placed in a darkened room and allowed to habituate for at least 1 hour.

For experiments involving optogenetic stimulation the light source was connected to a fiberoptic cable via a rotating commutator (FRJ_1×1_FC-FC, Doric Lenses) to allow freedom of motion. The fiberoptic cable was attached to the implant with a ferrule sleeve (Thorlabs) and mice were allowed to acclimate to the attached cable for 30 minutes. All assays were conducted under dim red illumination. Sensorimotor coordination was assessed with an automated rotarod apparatus (UgoBasile). Mice were placed on a rotarod with a constant rotation of 4 RPM, and allowed to acclimate for 1 minute, after which the rod accelerated to 60 RPM at a rate of 20 RPM/min. Time to fall was calculated from the beginning of acceleration. All mice were run on two consecutive trials with 4 minutes rest between trials, and animals were randomly assigned to receive optical stimulation during either the first or second trial. Optical stimulation began 10 seconds before the onset of acceleration and continued until the animal fell. Open field assays were conducted in a square opaque white plastic container (46 X 46 cm), and the central regions was defined as a square one third the dimension of the area. Automated tracking was performed in Matlab using idTracker2.1.

For resident-intruder assays, residents were attached to the optical fiber and allowed to acclimate in their home change for 30 minutes. A BALB/c intruder (roughly age matched) was introduced into the home cage, and interactions were filmed and manually scored. Trials were stopped in the event that either animal was injured by an attack, or if the resident attacked continuously for more than 60 s. Residents were run on up to 5 resident-intruder assays with at least 2 days between assays, with a novel intruder used for each assay. Residents were removed from the study if they failed to attack, or if the intruder attacked (Leshner & Nock, 1976). To establish a baseline level of aggression, assays with less than 3 attacks or more than 20 attacks were omitted from analysis. A subset of halorhodopsin-expressing animals (4/15) were stimulated in 5 minute intervals rather than the standard 1-minute intervals. Resident intruder assays were scored manually by an experimenter blinded to mouse genotype and stimulation wavelengths, and annotated using the open-source software BORIS (Friard & Gamba, 2016). The following behaviors were scored; self-grooming by the resident, social interactions (including face-to-face contact, mutual grooming and ano-genital sniffing), tail-rattles, lateral threat, chasing of the intruder by the resident, and biting attacks (Koolhaas *et al*., 2013).

## Acknowledgments

This work was supported by the G.V.R. Khodadad Research Fund for the Study of Excessive (Pathological) Selfishness and Aggressive Behavior. We especially wish to thank Ghahreman Khodadad for both his financial support and stimulating discussions. We also thank Jasmine Vazquez for illustrations, and Michelle Ocana and the Neurobiology Imaging Center for help with microscopy. This facility is supported in part by the Neural Imaging Center as part of a NINDS P30 Core Center grant (NS072030). This work was supported by a Nancy Lurie Marks postdoctoral fellowship to S.L.J., a NIH postdoctoral fellowship F32NS101889 to C.H.C, and NIH grant NINDS R35NS097284 to W.G.R.

---

**Figure 1 – figure supplement 1.**
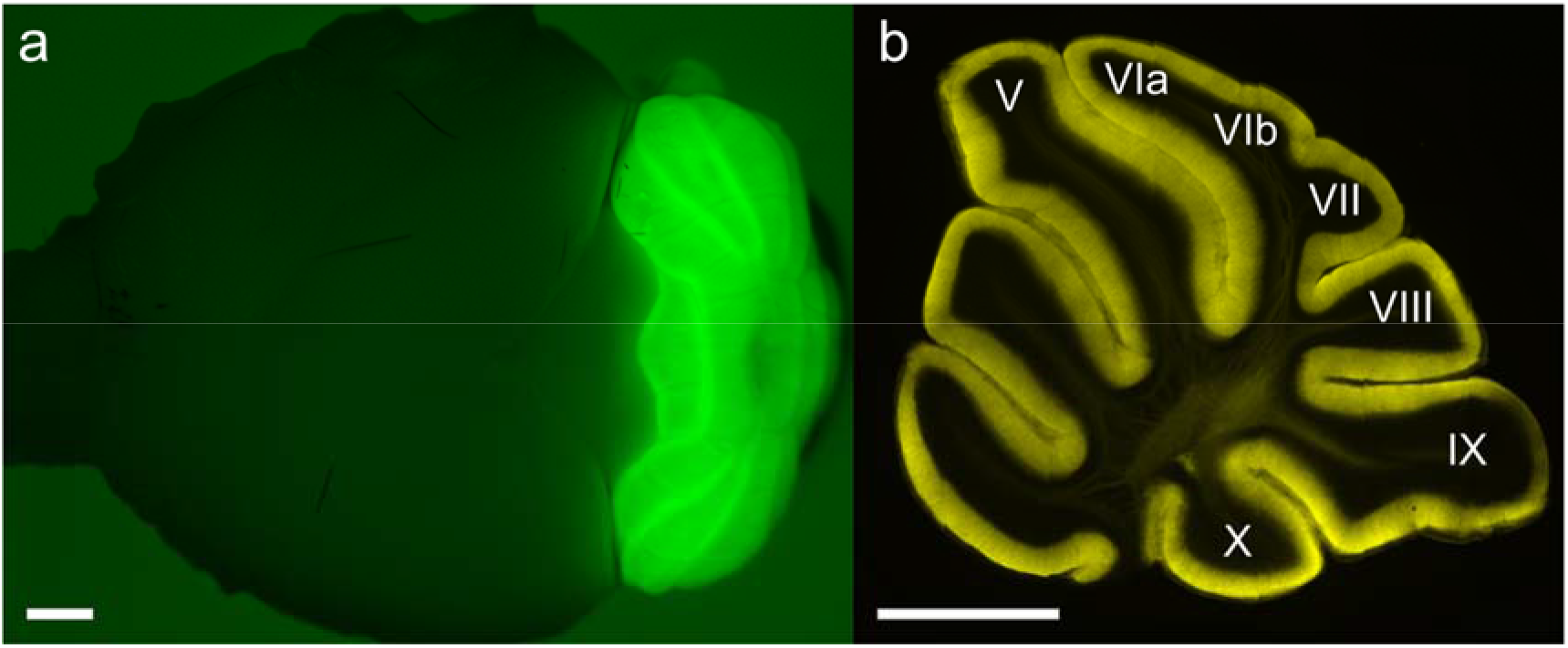
Fluorescent images of ChR2-YFP expression in (**a**) a whole brain and (**b**) a sagittal cerebellar section from a PCP2-cre::Ai32 mouse, with lobules V-X labeled. Scale bar = 1 mm.

**Figure 2 – figure supplement 1.**
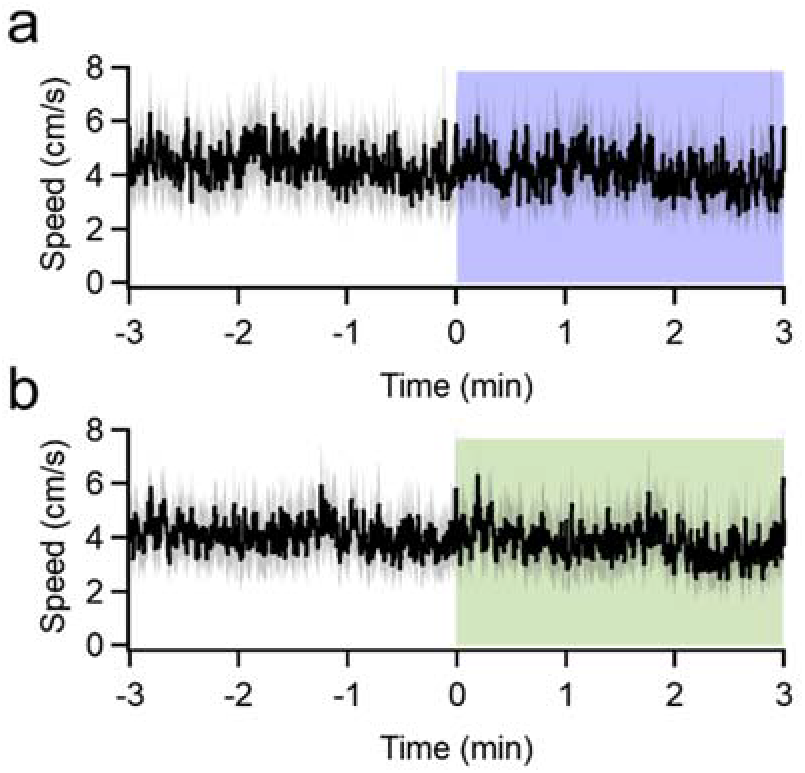
Manipulating vermal Purkinje cell firing does not affect locomotion. Animal speed in an open field, averaged across 3 consecutive epochs of stimulation for (**a**) PCP2-cre:ChR2 (n = 13) and (**b**) PCP2-cre:Halorhodopsin (n = 17) animals.

**Figure 3 – figure supplement 1.**
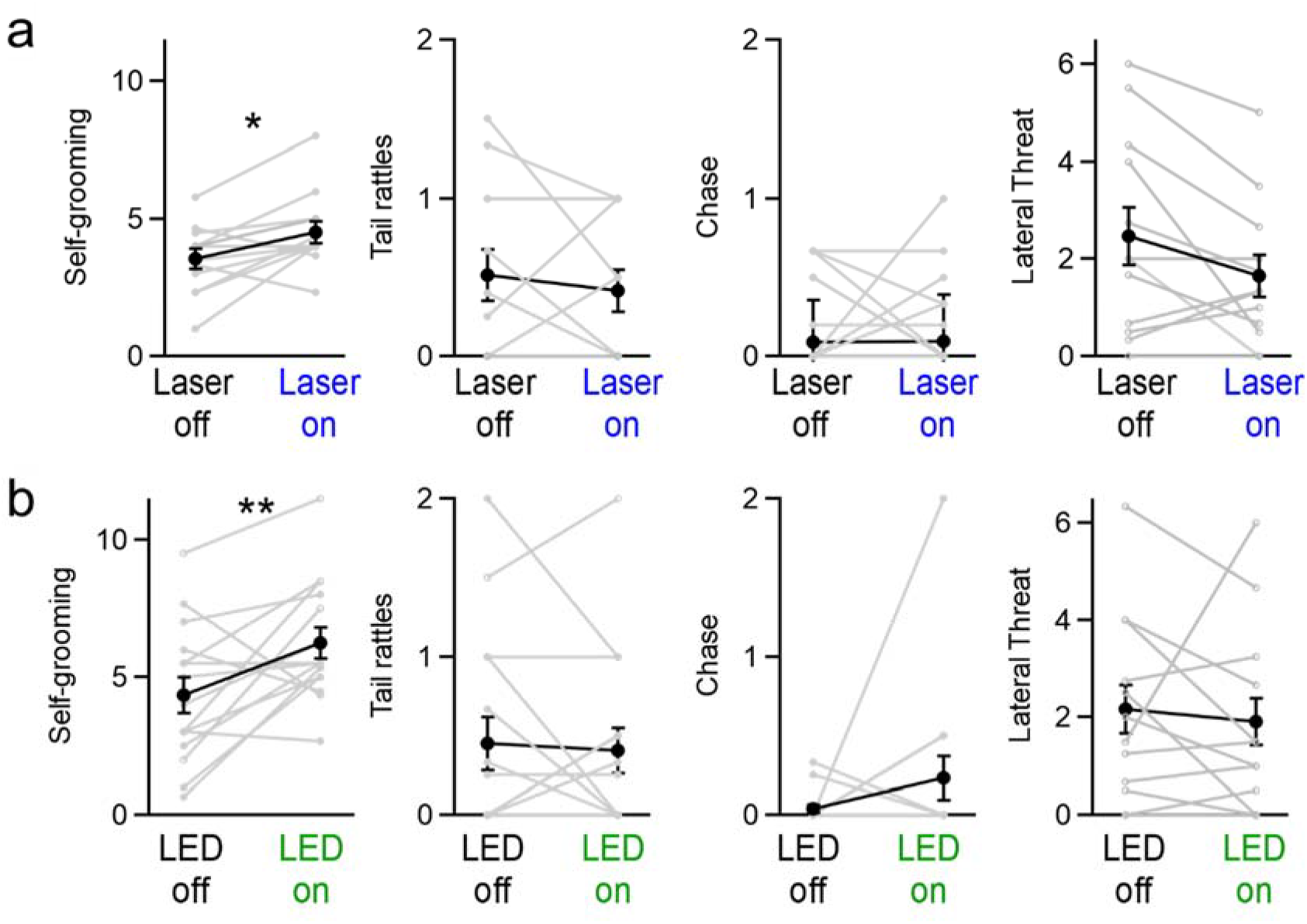
Effect of manipulating vermal Purkinje cell activity on grooming, tail-rattling, and aggressive lunging during resident-intruder assays. (**a**) Optogenetic stimulation significantly increased grooming behaviors for both ChR2-expressing mice (31 assays in 12 mice, *p* = 0.02, *t*-test) and (**b**) halorhodopsin-expressing mice (34 assays in 15 mice, *p* = 0.01, *t*-test).

**Figure 3 – figure supplement 2.**
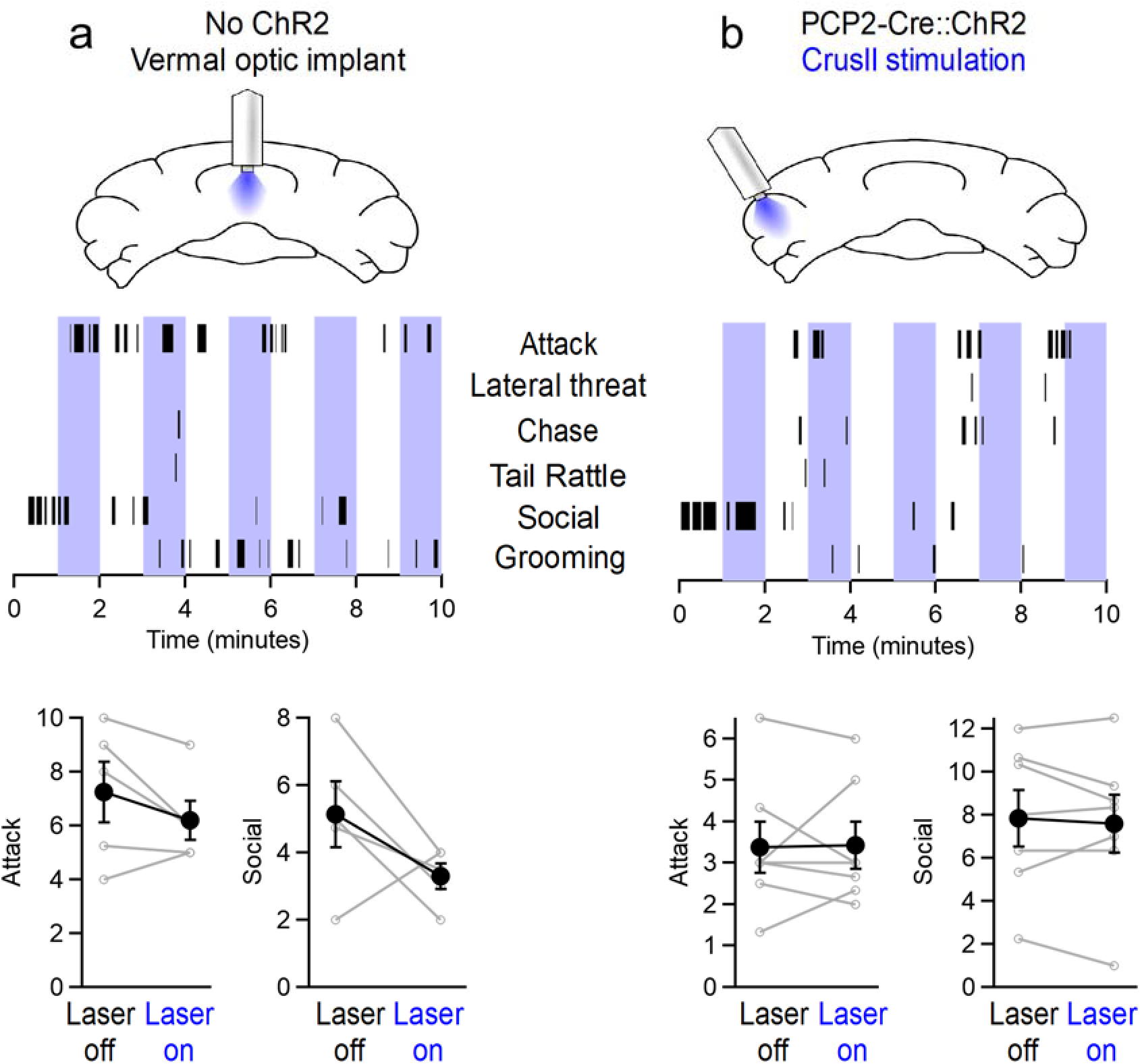
Aggression is not affected by light alone or by stimulating CrusII Purkinje cells. (**a**) (Top) Schematic for optical stimulation over vermis in wildtype mice lacking channelrhodopsin. (Middle) Representative sequence of social and aggressive behaviors. (Bottom) Average number of attacks and social investigations during ON/OFF epochs of laser stimulation. (**b**) Same as in (a), but for PCP2-Cre::ChR2-YFP mice implanted with optical fibers over the lateral CrusII region of the cerebellum.

